# Context-based protein function prediction in bacterial genomes

**DOI:** 10.1101/2024.10.14.618363

**Authors:** Daulet Toibazar, Maxat Kulmanov, Robert Hoehndorf

## Abstract

**Motivation:** The rapid growth of sequencing data from high-throughput technologies has emphasized the need to uncover the functions of unannotated genes. Recent advancements in deep learning algorithms have enabled researchers to utilize various features to predict protein functions. Traditionally, these algorithms treat proteins as independent functional units or consider interactions only at the protein level. However, prokaryotes often preserve specific genomic neighborhoods over evolutionary time, providing valuable context for predicting protein functions. This context can arise from genes near the gene of interest or synteny regions, where the conserved order of genes on chromosomes results from common ancestry.

**Results:** We developed a transformer-based model to pre-train representations of proteins based on their genomic context, and use this model for predicting protein functions. Our results show that context-based protein representations capture context-specific functional semantics and can effectively predict protein functions. We use our model to investigate the influence of phylogenetic distance and homology on the performance of context-dependent function prediction, and find that synteny affects the prediction performance substantially, except for some functions where the function is determined by the genomic context. Our experiments allow us to gain insights into the factors affecting the performance and applicability of context-based function prediction methods across diverse prokaryotic genomes and meta-genomes.

**Availability and implementation:** The generated model, including all training code and generated data, is freely available at https://github.com/bio-ontology-research-group/Genomic_context.

## Introduction

The rapid increase in genome and metagenome assemblies from diverse sources, driven by advancements in sequencing technologies and assembly methods have generated large amounts of genome and protein sequence data (Bateman *et al*., 2022; Delmont *et al*., 2011; Parks *et al*., 2017). Classical experimental methods to determine the function of proteins are highly informative (Markwick *et al*., 2008; Kemp and Alcock, 2017) but require substantial investment in terms of both time and materials, making them less accessible for large-scale or high-throughput studies. As a result, experimental function discovery has not kept pace with the rapid generation of new protein sequences, leading to a pronounced disparity between experimentally annotated and unannotated proteins.

The sequence–structure–function paradigm states that the function of a protein is largely dependent on its structure, which in turn is dependent on the protein sequence. While state-of-the-art models like AlphaFold3 (Abramson *et al*., 2024) and ESMFold (Lin *et al*., 2023) have largely solved the challenge of predicting protein structure from sequence, predicting function from sequence or structure remains an open research question. Computational approaches leveraging deep learning algorithms have been employed to tackle this issue, using diverse input features such as protein sequence (Kulmanov and Hoehndorf, 2019; Elhaj-Abdou *et al*., 2021; Kulmanov *et al*., 2021), protein–protein interaction (PPI) networks (Nguyen *et al*., 2011; Hou, 2017; Wan *et al*., 2019), protein structure (Konc *et al*., 2013; Lai and Xu, 2021), or multiple modalities (Cai *et al*., 2020; You *et al*., 2021). These methods show promising results in function prediction. However, they often treat proteins as independent functional units or only consider protein-level interactions.

The arrangement of genes, in particular in prokaryotic genomes, is not random; co-regulated genes are assembled in operons for more effective gene regulation, and conserved genome regions (syntenic blocks) are preserved under selective pressure (Rogozin, 2002; Guerrero *et al*., 2005; Junier and Rivoire, 2013; Osbourn and Field, 2009). The information present in a gene’s genomic context has been used for protein function prediction in some of the first computational function prediction methods (Huynen *et al*., 2000; Wolf *et al*., 2001). Novel machine learning algorithms are able to leverage genomic context information for discovering novel protein functions and novel protein interactions (Miller *et al*., 2022; Hwang *et al*., 2024). The word2vec model (Mikolov *et al*., 2013), particularly, has been used for context-dependent function prediction (Miller *et al*., 2022) with a comparable performance to sequence-based baseline models for selected functions. However, the word2vec model aggregates different genomic context variations, resulting in a static representation of genes and overlooking alternative functions or participation of genes in distinct biological pathways, depending on their genomic context. A more recent study (Hwang *et al*., 2024) leverages the Bidirectional Encoder Representations from Transformers (BERT) (Devlin *et al*., 2018) to contextualize sequence-based Evolutionary Scale Modelling 2 (ESM2) (Lin *et al*., 2023) embeddings and find that contextualized protein embeddings encode biologically relevant information that relates to protein function and their interactions. However, machine learning models trained on genomic context will be affected by synteny between genomes and may memorize synteny blocks even in the absence of protein sequence similarity to effectively predict protein functions. Furthermore, previous models based on word2vec or transformers only evaluated a small set of manually selected functions based on prior hypotheses about which functions may be dependent on genomic context.

To overcome these limitations, we developed a novel transformer-based model that exploits genomic context for protein function prediction. We test our model using InterPro IDs and a selected set of GO functions. We further evaluate the effects of phylogenetic relatedness and synteny on the performance of our context-based protein function prediction using our model. We demonstrate that our model can improve over prior models for context-based function prediction. Our model, including all training code and generated data, is freely available at https://github.com/bio-ontology-research-group/Genomic_context.

## Materials and methods

### Prokaryotic genome assemblies and gene clustering

We downloaded a genome dataset from NCBI, available on January 30, 2023, (https://www.ncbi.nlm.nih.gov/assembly). The dataset comprised a total of 31,002 genomes containing 119,851,800 protein sequences. We clustered the 119,851,800 (putative) protein sequences using the MMseqs2 (Steinegger and Söding, 2017) linclust algorithm, setting the `min-seq-id` parameter to 0.3. Clustering resulted in 8,988,233 clusters. We removed clusters with fewer than 20 sequences to further refine the dataset, eliminating approximately 18% of the original protein sequences. This filtration step reduced the number of clusters to 544,993 while retaining most initial sequences. We assigned a representative protein sequence to each cluster using mmseqs createsubdb DB_clu DB_DB_clu rep command. We used the InterProScan version 5.66-98.0 (Jones *et al*., 2014) to assign InterPro IDs to each cluster representative protein. These InterPro IDs serve as ground-truth labels for their corresponding protein sequences in further analysis.

### Reformulating genome corpus and creating MLM dataset

Following Miller *et al*. (2022), we reformulated the genome corpus by substituting protein sequences with cluster-representative protein accession IDs. Replacing the protein sequences with protein IDs transforms the genomes into a “corpus” where tokens are proteins.

We iterated through the genome corpus to generate the Masked Language Modeling (MLM) pre-training dataset, where all sentences consisted of 9 proteins. We divided the dataset into two parts: 90% for training and 10% for evaluation; we used the validation set to monitor the model’s training process and employed early stopping if the evaluation loss ceased to improve.

### BERT architecture and pre-training

We developed a custom tokenizer using the HuggingFace (Wolf *et al*., 2019) tokenizer library with a vocabulary size of 544,998 (one for each cluster representative protein) to ensure each protein cluster is recognized as a separate token.

We chose BERT as the base model for this project due to its bidirectional encoding strategy, which enables the comprehension of contextual information from input sentences. We used BERT with the following parameters: `number hidden layers = 12, hidden size = 512, num attention heads = 8`. We pre-trained the BERT model *de novo* using the self-supervised MLM objective, where we randomly masked 20% of the input tokens, and the model predicts masked tokens based on the remaining tokens in the input sentence.

The pre-training process used 8 V100 GPUs to accelerate the computations and enable efficient training on the large dataset. To streamline the training process, we used the HuggingFace (Wolf *et al*., 2019) `Trainer` class. This class provides a high-level interface for training models, handling data loading, and managing the training loop.

After pre-training, we evaluated the BERT with the “Fill-Mask” task, where we assessed the BERT’s ability to correctly predict the masked gene in a chunk of the genome using top-1 and top-5 accuracy. Top-1 accuracy takes into consideration the token with the highest predicted probability 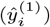 and compares it to ground truth, while top-5 accuracy takes into consideration the 5 highest prediction probability tokens 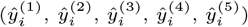.

### Extracting BERT embeddings and functional annotation

We used our pre-trained BERT model to extract contextual protein representations by passing chunks of the genome: **E** = BERT(**I**) (Supplementary Figure 1). Let **I** be the input to the model with **n** sentences and *l* representing the sentence length, **I** *∈* ℝ^**n***×l*^.

From protein embeddings, denoted as **E** *∈* ℝ^*n×l×h*^, where *h* represents the embedding size, we extracted the embeddings for the proteins in the middle of each sentence. We annotated the middle protein to ensure balanced context-aware embeddings from both sides. Using proteins from different positions can yield varying embeddings due to slight shifts in the context window. Averaging embeddings in a sentence is redundant since each token embedding already contains contextual information about surrounding tokens. We obtained the corresponding labels for each **x**_*k*_ *∈* ℝ^*h*^, *k ∈* [1, 2, …, *n*], using a multilabel binarized function from the *sklearn* library (Pedregosa *et al*., 2011). We presented the labels in a one-hot encoded form in a matrix *Y ∈*ℝ ^*n×c*^, where we define *C* as the set of all InterPro classes and |*C*| = *c*. In matrix Y, a value of 1 indicates that the given gene is annotated with a certain InterPro ID, and 0 is otherwise.

For the supervised classification task, we trained a prediction MLP model with 3 hidden layers, configured with weights *W*_*i*_ and biases *b*_*i*_, where *i ∈* [0, 1, 2, 3]. The input layer accepts the embedding matrix *X ∈*ℝ^*n×h*^, and the output layer produces *Ŷ ∈*ℝ ^*n×c*^.

We used BCEWithLogitsLoss (Paszke *et al*., 2019) as the loss function to enable the multilabel classification objective. Additionally, we employed the Adam (Kingma and Ba, 2014) optimizer with a learning rate of 0.001, which decayed by a gamma factor of 0.05 every 5 epochs.

### Incorporating phylogeny information into dataset splits

To investigate the impact of evolutionary distance on contextual predictive performance, we divided the genome dataset into various train/test splits based on phylogeny (de Vienne, 2016). Besides a **random** split, where all genomes were randomly split into train and test sets, we utilized splits at the **family** (e.g., *Enterobacteriaceae*), **order** (e.g., *Enterobacterales*), and **class** (e.g., *Gammaproteobacteria*) levels. In each split, all genomes from a particular family were allocated to the test set, while the rest of the genomes outside of the specified phylogenetic circles were used to train the model (Supplementary Figure 2). We systematically widened the evolutionary gap between training and testing datasets to evaluate the model’s ability to capture and leverage genomic contextual cues beyond those confined to synteny regions or operons inherited from shared ancestors. We extracted BERT embeddings for all splits and trained an MLP classifier to annotate their corresponding InterPro IDs. We repeated this experiment using ESM2 embeddings derived from cluster-representative protein sequences to analyze the effect of evolutionary distance on sequence-based models and establish a baseline for our contextual approach. Specifically, we utilized the *esm2 t30 150M UR50D* model, featuring 30 layers and 640 embedding dimensions.

### Evaluation metrics

We assess our experiments with two evaluation metrics: *F*_max_ and AUC-ROC scores. The first metric is a protein-centric maximum value for the F score with threshold, *t ∈* [0, 1]. We used the standard *F*_max_ formulation used in the CAFA (Radivojac *et al*., 2013) challenge, where we define *I*_*t*_ as a set of proteins annotated with at least one InterPro (Eq. 1). Precision is averaged over proteins in *I*_*t*_ (Eq. 2), while recall is averaged over all proteins in the dataset (Eq. 3). Then, *F*_max_ is chosen from the maximum value of harmonic precision-recall mean over all possible values of *t* (Eq. 4).

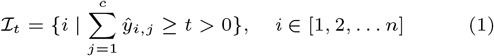

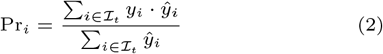

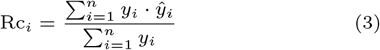

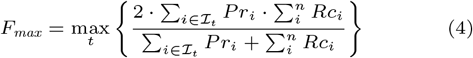

The second evaluation metric is an InterPro term-centric AUC for the Receiver Operating Characteristic (ROC) curve. ROC curve shows the performance of our model at all classification thresholds. It plots the true-positive rate (TPR) against the false-positive rate (FPR). We calculate TPR and FPR for class *j, j ∈* [1, 2, …, *c*] with the following equations:

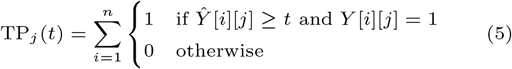

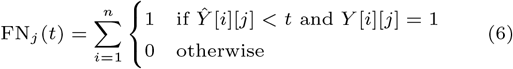

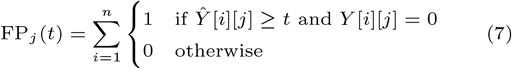

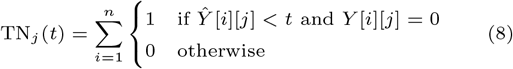

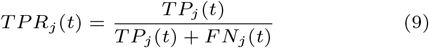

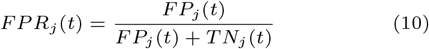

Then, using TPR and FPR, we integrate over threshold values to find the AUC for the ROC curve. The final AUC value is calculated by taking *micro* and *macro* average of class-centric AUC values.

### Statistical analysis

To assess the statistical significance of differences in *F*_max_ values across various train/test splits, we conducted 4 trials for each experiment. Each trial used different random seeds for MLP model weight initialization and data shuffling, allowing us to add Gaussian noise to our experiments. We then used a one-way ANOVA (Fisher, 1925) (Analysis of Variance) test to determine if there are statistically significant differences in the averaged *F*_max_ between random, family, order, and class-level splits.

If the ANOVA indicated significant differences between the groups, we conducted a post-hoc analysis using Tukey’s Honest Significant Difference (HSD) (Tukey, 1949) test to identify which specific splits were statistically different.

### Extending function prediction to Gene Ontology

We extended our protein function prediction task by incorporating experimental protein annotations from QuickGO (https://www.ebi.ac.uk/QuickGO/annotations) across three Gene Ontology (GO) branches: Molecular Function (MF), Biological Process (BP), and Cellular Component (CC). To annotate proteins in our corpus, we performed BLAST searches against these experimentally annotated proteins, using a sequence similarity threshold of *>* 90%. After completing the annotation process, we extracted BERT embeddings from proteins with experimental GO annotations. These embeddings served as input for training an MLP model, which learned to map them to corresponding GO functions. Term-centric AUC scores provided a metric for evaluating our prediction effectiveness. To focus our analysis, the results were filtered to highlight the top-*n* highest AUC scores along with their associated GO terms. Contextualizing these findings involved a performance comparison between our model and the state-of-the-art DeepGO-SE (Kulmanov *et al*., 2023) model. This comparative analysis aimed to explore the potential of genomic context information, as captured by BERT embeddings, in complementing the features utilized by the DeepGO-SE model.

### Investigating the generalization of BERT to diverse genomic architectures in prokaryotes

Further, we compared our BERT model against the word2vec to investigate which model generalizes better to the diverse genomic contexts in bacterial genomes. For this comparison, we utilized the corpus proposed by Miller et al. (Miller *et al*., 2022), where they provide gene KEGG Orthology (KO) identifiers and categorize genes into functional classes. From all functional classes available in metadata, we selected the same 9 classes used in the original paper (Miller *et al*., 2022), excluding the ‘Others’ class. We extract all KO’s corresponding to 9 classes and obtain their context-dependent embeddings.

For the BERT model, we pre-trained BERT using the MLM objective on the published genome corpus (Miller *et al*., 2022) and applied a similar strategy as shown in Supplementary Figure 1. To evaluate the word2vec model, we average the embeddings by considering 4 contextual proteins from both sides of the protein of interest, making the representation more context-specific (Supplementary Figure 3); otherwise, all embeddings from the same KO would be identical despite differences in the genomic context. Finally, we trained an MLP classifier that assigns a functional class to each embedding and evaluate the classification task using precision, recall, and F1 scores. We performed an evaluation with 3 repetitions, each time sampling 10,000 genomes from the corpus.

## Results

### Genomic language model: pre-training and evaluation

We developed an encoder-based BERT model to assess if protein representation learning from genomic context suffices for protein function prediction in prokaryotes. Our method involved creating a consistent vocabulary across genomes by clustering proteins at 30% sequence identity and replacing proteins with their cluster’s accession identifier. We trained a BERT model using a masked language modeling objective, where 20% of the tokens are masked, and the model predicts them based on their genomic context. The objective of the training was to learn meaningful representations of proteins that capture their contextual relationships within the genomic regions where they appear. We conducted a “Fill-Mask” evaluation to assess the performance of the pre-trained BERT model. This method involves predicting a masked token within a sequence of tokens, allowing us to measure the model’s accuracy in identifying the correct token. The evaluation revealed a 94% top-1 accuracy and a 99% top-5 accuracy across 500 randomly selected genomes.

Next, we predict protein functions using the proteins’ BERT embeddings and a prediction model. We divide our evaluation into three groups: general proteins (including proteins with enzymatic functions, such as synthases, dehydrogenases, ATPases, etc.), secretion proteins, and defense system proteins. We use the proteins’ InterPro identifiers as functional labels for each protein and design a prediction model to predict these functional labels from the BERT embeddings. The prediction model is represented by a multi-layer perception (MLP) model with 3 hidden layers and a ReLU activation function. Supplementary Figure 1 illustrates the workflow and how we perform gene function prediction. We benchmark context-dependent BERT embeddings against sequence-based ESM2 embeddings to assess the extent of genomic context information compared to protein sequence information for listed classes of bacterial proteins. Our results (Table 1) show that defense system proteins achieve the highest *F*_max_ value when predicted from BERT embeddings, followed by general and secretion proteins. On the other hand, predictions based on ESM2 embeddings achieve the highest *F*_max_ for the general protein class, followed by secretion and defense proteins. Our findings show that, on average, the ESM-based model that uses protein sequences as input can better predict general protein function than it can predict functions related to the defense or secretion system, which are inherently context-dependent functions; the BERT-based model which exploits genomic context shows the opposite trend. Furthermore, we also find that BERT embeddings achieve higher *F*_max_ and AUC values for all three types of proteins compared to ESM2 embeddings.

**Table 1.** Context-based function prediction performance in comparison to sequence-based ESM2 embeddings.

|  | BERT |  |  | ESM2 |  |  |
| --- | --- | --- | --- | --- | --- | --- |
|  | defense | secretion | general | defense | secretion | general |
| $F_{\max}$ | 0.659 | 0.595 | 0.596 | 0.136 | 0.162 | 0.214 |
| AUC | 0.968 | 0.966 | 0.950 | 0.829 | 0.880 | 0.937 |

### Evaluating the impact of phylogenetic distance on context-based function prediction performance

We hypothesize that the increased performance observed for the context-based model comes from a combination of two phenomena: (1) functions that are inherently dependent on the genomic context (such as bacterial defense and secretion), and which the BERT model can exploit; and (2) synteny regions appearing in closely related organisms and which our MLP model memorizes when it sees the related genome during training.

To test the extent to which defense and general protein classes’ genomic context information is influenced by synteny in closely related organisms, we designed an experiment to sequentially reduce the impact of synteny. In our experiment, we split the training and testing sets (consisting of entire genomes) according to their phylogenetic origin. Splitting organisms according to phylogenetic origin increases the evolutionary distance between the genomes in training and testing, and limits the ability of the MLP model to memorize genomic context. We systematically increase the phylogenetic distance from a random split to a split by family, order, and class levels. The bacterial families we analyzed include *Enterobacteriaceae* and *Pseudomonadaceae*, as these are the largest families in our genome dataset. At each phylogenetic level, we use genomes within the phylogenetic group as the test set; all genomes outside of the specified phylogenetic group are used as the training set (Supplementary Figure 2).

The results of our experiments (Table 2) demonstrate a substantial drop in performance as we increase the phylogenetic distance between training and testing sets. The drop in predictive performance from a random split to family, order, and class are all significant (*p ≈* 0.0); there is also a significant drop between order- and class-level splits (*p* = 0.019). For general proteins, there is a significant difference in *F*_max_ between random and any phylogeny-based split (*p ≈* 0.0) and between the family–class (*p* = 0.002) and family–order (*p* = 0.040) level splits. Supplementary Tables 1 and 2 contain detailed results of statistical analysis.

**Table 2.** Comparative analysis of *F*_max_ scores for defense and general class proteins using BERT embeddings across phylogenetic distance.

|  | Defense |  | General |  |
| --- | --- | --- | --- | --- |
|  | Mean | Std | Mean | Std |
| Random | 0.659 | 0.025 | 0.596 | 0.008 |
| Family | 0.356 | 0.039 | 0.318 | 0.038 |
| Order | 0.342 | 0.053 | 0.271 | 0.034 |
| Class | 0.281 | 0.022 | 0.249 | 0.037 |

Our results further demonstrate that defense proteins (which are known to be context-specific) achieve higher *F*_max_ values in all phylogenetic splits when compared to general proteins (Supplementary Figure 4).

### Comparative study of phylogenetic distance on sequence-based models

Models that use protein sequences should be less sensitive to increasing the evolutionary distance between training and testing, and should also be more accurate for general proteins. The hypothesis is based on the assumption that context-dependent systems, such as defense systems, often rely on genetic organization along with protein sequence for proper functioning (Makarova *et al*., 2011; Doron *et al*., 2018). In contrast, general protein classes are expected to be less dependent (or independent) on contextual cues and more reliant on the intrinsic properties of the proteins themselves.

Therefore, we repeat the experiment using the ESM2-based model (Table 3). We observe that, as we increase the evolutionary distances from family over order to class for both *Enterobacteriaceae* and *Pseudomonadaceae* families, we observe a significant difference for defense proteins only when transitioning directly from a random split to a class-level split (*p* = 0.036). A small decrease in mean *F*_max_ values for defense proteins can likely be attributed to a higher protein sequence similarity between closely related organisms, and as we increase evolutionary distance, we decrease sequence similarity between the training and test sets for the MLP model. For general proteins, we observe no significant difference between any splits (*p ≥* 0.05). Furthermore, compared to defense proteins, general proteins achieve a higher *F*_max_ across all splits. This observation aligns with our initial hypothesis that general proteins (e.g., enzymes) depend mainly on the protein’s intrinsic factors (i.e., sequence and structure) rather than genomic context.

**Table 3.** Comparative analysis of *F*_max_ scores for defense and general class proteins using ESM2 embeddings across phylogenetic distance.

|  | Defense |  | General |  |
| --- | --- | --- | --- | --- |
|  | Mean | Std | Mean | Std |
| Random | 0.136 | 0.028 | 0.214 | 0.070 |
| Family | 0.116 | 0.031 | 0.140 | 0.038 |
| Order | 0.096 | 0.023 | 0.146 | 0.050 |
| Class | 0.088 | 0.019 | 0.156 | 0.043 |

### Context-only dependent Gene Ontology function prediction

We extracted BERT embeddings from proteins with experimental GO annotations and trained an MLP model to predict GO functions from the embeddings. Our model’s performance was evaluated using term-centric AUC scores and compared against the state-of-the-art DeepGO-SE model. The terms that achieved the highest AUC scores from our BERT-based approach mostly showed AUC values close to 0.5 in DeepGO-SE predictions. Specifically: 17 out of 20 terms achieved higher AUC scores in BP, 15 out of 20 terms showed improved AUC values in MF, and 12 out of 15 terms exhibited better AUC scores in CC. The remaining terms performed similarly or slightly better with DeepGO-SE. Detailed results for each GO aspect are provided in Supplementary Tables 5, 6, and 7.

One example of a function that our genomic context model can predict but DeepGO-SE cannot is “plasmid recombination” (GO:0042150). Plasmid recombination is influenced significantly by the genomic context, specifically the genes flanking the recombination sites on both the plasmid and the host genome. Homology between these flanking genes enhances the likelihood of homologous recombination, as sequence similarity provides the necessary substrate for the recombination machinery to align and exchange genetic material accurately (mol, 2022). Additionally, the presence of repetitive gene sequences, such as those generated by transposons, can drive non-homologous recombination, further emphasizing the role of surrounding genes in the process (mol, 2022). Given this dependency of this function on surrounding genes, it is plausible that the prediction of plasmid recombination as a protein function can be achieved more effectively from the genomic context than from sequence as done by DeepGO-SE.

Our findings indicate that genomic context, as captured by BERT embeddings, provides valuable information for protein function prediction. This contextual information can be complementary to sequence-based information and therefore enhance the predictive power of existing models like DeepGO-SE.

### BERT generalizes better for different genomic contexts in bacterial genomes

Our model is based on transformers; however, we could also have used (contextualized) word embeddings using a model such as word2vec instead. To determine whether the transformer model has advantages over word2vec, we pre-trained our BERT model on a corpus published by Miller et al. (Miller *et al*., 2022) and performed multi-class classification task using 9 classes presented in the original paper, except for the “Others” class. We also modified the word2vec classification pipeline to generate context-specific embeddings to ensure a fair comparison with BERT: we averaged word2vec embeddings across 9 genes, having 4 contextual genes from both sides. The results in Table 4 show that the transformer-based language model outperforms word2vec embeddings in *F*_max_ score (*p* = 4.652 *·* 10^*−*9^). The BERT model, and its attention heads, learn a more comprehensive and context-specific functional representations of genes and are better at leveraging contextual information within bacterial genomes than the contextualized word2vec model. The performance metrics per class for word2vec and BERT are given in Supplementary Tables 3 and 4, respectively.

**Table 4.**
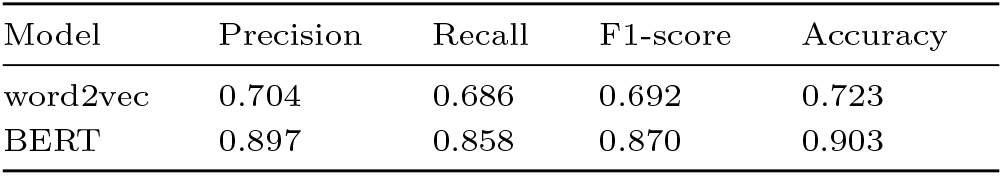
Functional classification results for BERT and contextualized word2vec embeddings.

| Model | Precision | Recall | F1-score | Accuracy |
| --- | --- | --- | --- | --- |
| word2vec | 0.704 | 0.686 | 0.692 | 0.723 |
| BERT | 0.897 | 0.858 | 0.870 | 0.903 |

## Discussion

### Effects of evolutionary distance and synteny

Our study shows that protein functions can be predicted using only their genomic context as input, substantially improving predictive performance over models that use protein sequence (or their corresponding ESM2 embedding) as input at least for the classes of protein functions we evaluated (i.e., functions related to defense and secretion). Our findings also align with prior work that used word2vec (Miller *et al*., 2022) to predict these functions or that combined ESM2 embeddings with genomic context to form contextualized protein embeddings (Hwang *et al*., 2024). However, in contrast to prior work, we have developed a model that uses only the genomic context.

Using our genomic context-based model, we were able to identify two distinct sources of information that are used to predict the protein functions: the genomic context, and evolutionary relatedness and synteny between genomes used in training and testing. The first aligns with our hypotheses and also the conclusions of prior work that finds genomic context to be important in predicting some protein functions; the second source of information, however, directly contradicts these conclusions. Exploiting similar or identical genomic arrangements between training and testing genomes (i.e., synteny) is usually undesirable as it severely limits the model’s ability to generalize, and results obtained by exploiting this information amount to a kind of “overfitting” of the model where it can perform well by only memorizing genomic arrangements seen during training.

Therefore, we designed an experiment to evaluate the different protein categories under different phylogenetic distances – from random splits to family, order, and class-level splits; our experiment revealed that the genomic context models indeed exploit synteny. Synteny is due either to inheritance of the same regions from common ancestors or, in bacterial genomes, acquired through horizontal gene transfer. Our experiments provide an avenue to remove the effect of synteny from the prediction models. Such an approach is common in sequence-based function prediction where training proteins are usually expected to be sufficiently distinct from testing proteins (e.g., by employing a split based on sequence similarity).

We also show that different functions are affected differently by synteny as well as by genomic context. Bacterial defense proteins can consistently be predicted better from genomic context than general proteins across all phylogenetic levels, indicating their functional predictability is less affected by phylogenetic distance. Furthermore, the prediction performance also declines significantly only when moving from random split to any phylogenetic split, whereas there is no significant difference when moving from family to order, or order to class. This finding supports our hypothesis that the predictive ability of contextual information is closely tied to conserved genomic regions and interactions with neighboring genes, i.e., that, when predicting defense mechanism functions, genomic context is more important than synteny, i.e., genomic context is what conveys the function to the proteins.

We observe the opposite for general proteins with a pronounced and significant decay in predictive performance as contextual information increases. This decay suggests that the ability to predict protein function in general proteins is more sensitive to phylogenetic divergence, i.e., the performance of the BERT model primarily comes from synteny and exploiting similarity to genomic context seen for closely related genomes during training.

### Transformer generalize word2vec models for context-based function prediction

Language models trained with self-supervised objectives have demonstrated state-of-the-art performance in many NLP tasks (Raiaan *et al*., 2024), proving their ability to understand language’s syntactic and semantic structure. In particular the attention mechanisms and the increased capacity of the model over word2vec representations can improve performance. In our study, the BERT model shows a superior performance compared to contextualized word2vec embeddings.

However, improved predictive performance is not the only advantage of the BERT-based architecture. While the word2vec model can predict protein function based on context, it will output the same function for a protein independent of its context; however, proteins, even if they are sequence identical, can be involved in different functions in different genomes and, therefore, in different genomic contexts. Recent studies have shown that the CRISPR-Cas system, initially known for its role in bacterial immunity, is involved in functions beyond this primary purpose (Devi *et al*., 2022). We conducted a case study on the branch family of Cas2 protein (IPR010152) to visualize the contextual embeddings this protein generates in different bacterial genomes. Our findings revealed that there is one cluster located far from the other clusters, suggesting a function beyond adaptive immunity (Supplementary Figure 5).

## Conclusion

We developed a model for protein function prediction based on genomic context. Our model is based on transformers and trained using a self-supervised masked language modeling objective. We demonstrate that our model improves over state-of-the-art models for context-based protein function predictions that are based on word2vec. Our model is also the first transformer model that is solely based on genomic context instead of combining genomic context with sequence-based representations. This separation allows us to demonstrate a kind of overfitting in context-based function prediction where the models memorize synteny between different genomes instead of actually learning the “semantics” of arrangements of genes in a genome and recognizing where this arrangement functions; it also allows us to remove or reduce this overfitting to synteny. Our model, together with all its training code, is freely available, allowing other researchers to build on and extend it.

### Limitations and future research

In the future, we aim to leverage the Gene Ontology (GO) or KEGG functions more substantially because they offer more informative functional categories; for example, GO annotations encompass biological processes, molecular functions, and cellular components, offering a broader and more granular view of protein functions.

To further advance this field of research, future studies focus on attention scores in different genome regions, which can open new insights into genetic interactions and operons. This approach would provide a more comprehensive understanding of the biological processes in which the gene is involved and the functional units of which they are a part. Another avenue for future research is the analysis of context-dependent function annotation and the effect of evolutionary relatedness in eukaryotic genomes, which have been reported to possess macro- and microsynteny structures (Li *et al*., 2022; Simakov *et al*., 2022). Investigating the impact of genomic context on protein function prediction in eukaryotes could reveal novel insights into the evolutionary conservation and divergence of functional modules across different domains of life.

## Author contributions statement

D.T. and R.H. conceived the experiments, D.T. conducted the experiments, D.T., M.K. and R.H. analyzed the results. D.T., R.H., and M.K. wrote and reviewed the manuscript. M.K. and R.H. obtained funding and supervised the research. All authors have read and approved the final version of the manuscript.

## Acknowledgments

This work has been supported by funding from King Abdullah University of Science and Technology (KAUST) Office of Sponsored Research (OSR) under Award No. URF/1/4675-01-01, URF/1/4697-01-01, URF/1/5041-01-01, REI/1/5659-01-01, REI/1/5235-01-01, and FCC/1/1976-46-01. We acknowledge support from the KAUST Supercomputing Laboratory.

