## Supplementary_data for "Context-based protein function prediction in bacterial genomes"

Supplementary Material

**Daulet Toibazar[1], Maxat Kulmanov[2], and Robert Hoehndorf[2]**

[1] Biological and Environmental Science and Engineering Division, King Abdullah University of Science and Technology, Thuwal, Kingdom of Saudi Arabia
[2] Computer, Electrical and Mathematical Sciences & Engineering Division, King Abdullah University of Science and Technology, Thuwal, Kingdom of Saudi Arabia


#### List of Figures

1 The protein function prediction pipeline operates as follows: (1) The BERT model inputs a genome segment, with each token as a distinct protein, generating a 512-dimensional embedding for each token. We use the embedding at index 4 for classification. (2) The MLP model takes this 512-dimensional embedding and outputs prediction probabilities for all $c$ InterPro classes. . . . . . . . . 2

2 Miniature phylogenetic tree illustrating training and testing set splits based on evolutionary distance. Each circle delineates a split level, where genomes of taxa located outside the circle are utilized for training the classification model. The blue and green circles denote two bacterial lineages we utilized in this research. As the circle size increases, the evolutionary distance between the training and testing sets becomes larger. Notably, across all split levels, only genomes from the *Enterobacteriaceae* and *Pseudomonadaceae* families are used for testing. This approach ensures a consistent distribution of genes in the test set, enabling a fair comparison of the model's performance at different evolutionary distances . . . . . . . . . 3

3 Overview of contextualizing word2vec embeddings and predicting corresponding functional class. The numbers inside boxes denote embedding dimensions. . . . . . . . . 4

4 Comparative analysis of Fmax scores for defense and general genes using BERT and ESM2 embeddings. Panel (a) displays the scores for defense genes, while panel (b) presents the scores for general genes. Each panel compares the performance of context-dependent BERT embeddings and sequence-based ESM2 embeddings across different train/test splits: Random, Family-level, Order-level, and Class-level. . . . . . . . . 5

5 Visualization of Cas2 protein embeddings, serving as a case study. The legend displays various protein identifiers, all annotated with the InterPro ID IPR010152, which corresponds to Cas2 proteins. The plot reveals two distinct clusters: a main cluster and a smaller, remote cluster. This separation suggests that some Cas2 proteins exist in different genomic contexts. The distinct clustering highlights the importance of considering genomic context when analyzing protein function, even within the same protein family. . . . . . . . . 6

7 Comparison of AUC Scores for BERT's Top 15 Cellular Component GO Terms vs. DeepGO-SE . . 10

### 1 Supplementary Figures

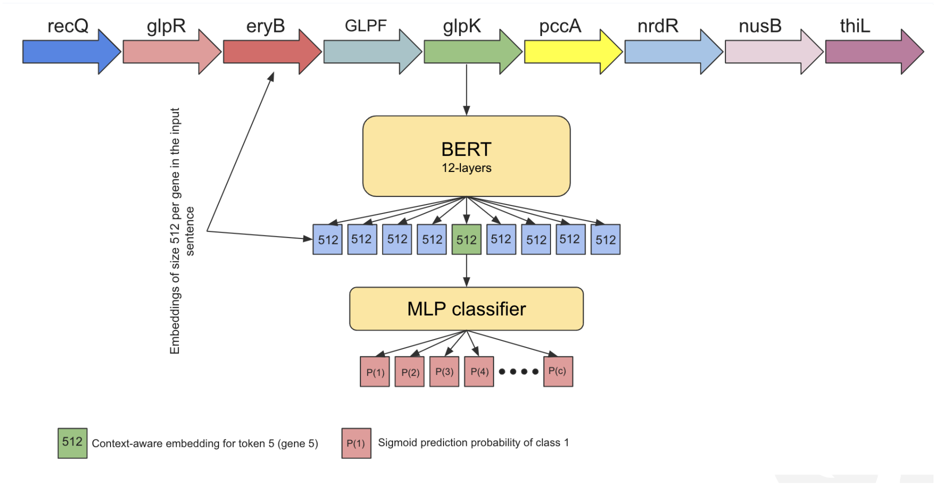

Figure 1: The protein function prediction pipeline operates as follows: (1) The BERT model inputs a genome segment, with each token as a distinct protein, generating a 512-dimensional embedding for each token. We use the embedding at index 4 for classification. (2) The MLP model takes this 512-dimensional embedding and outputs prediction probabilities for all $c$ InterPro classes.

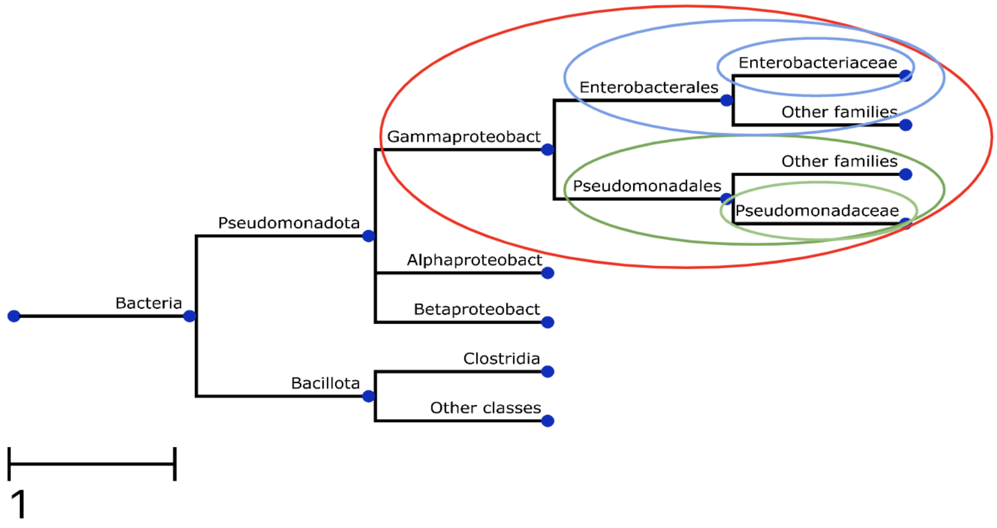

Figure 2: Miniature phylogenetic tree illustrating training and testing set splits based on evolutionary distance. Each circle delineates a split level, where genomes of taxa located outside the circle are utilized for training the classification model. The blue and green circles denote two bacterial lineages we utilized in this research. As the circle size increases, the evolutionary distance between the training and testing sets becomes larger. Notably, across all split levels, only genomes from the *Enterobacteriaceae* and *Pseudomonadaceae* families are used for testing. This approach ensures a consistent distribution of genes in the test set, enabling a fair comparison of the model's performance at different evolutionary distances

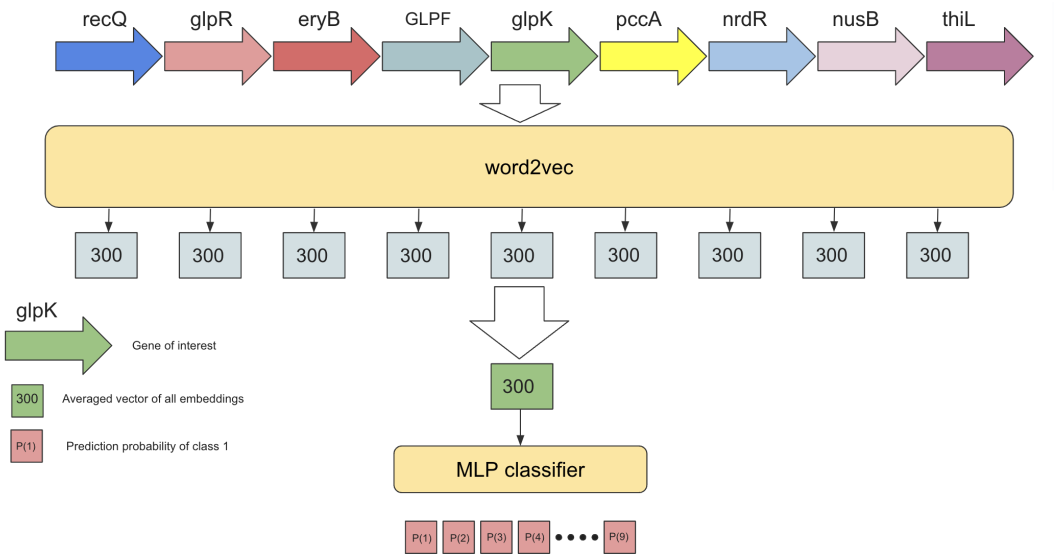

Figure 3: Overview of contextualizing word2vec embeddings and predicting corresponding functional class. The numbers inside boxes denote embedding dimensions.

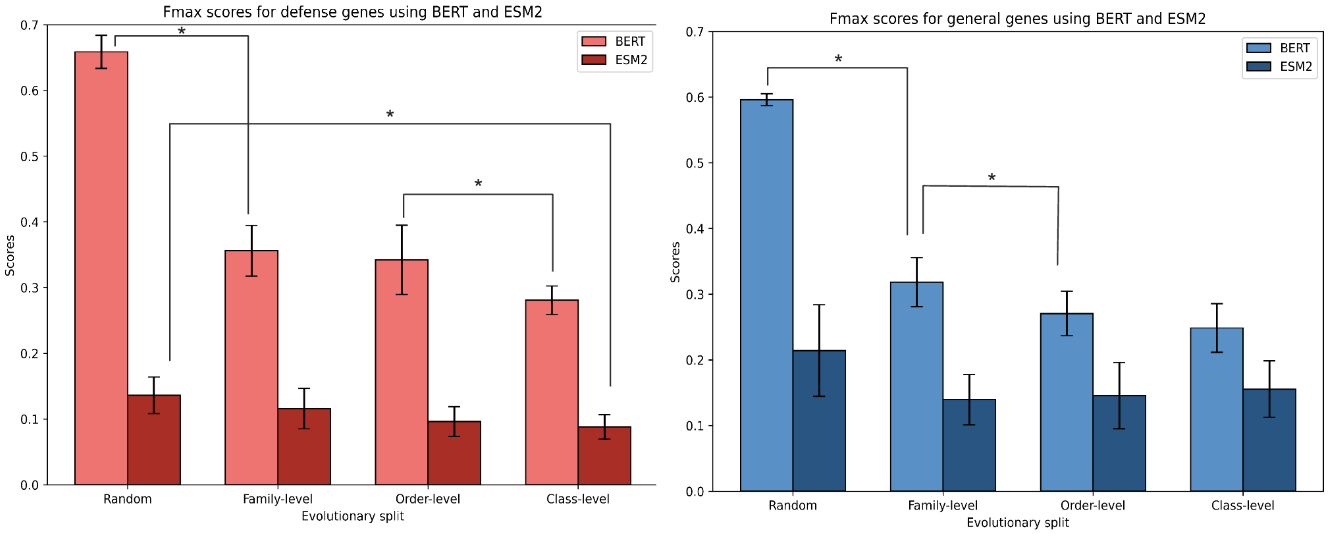

Figure 4: Comparative analysis of Fmax scores for defense and general genes using BERT and ESM2 embeddings. Panel (a) displays the scores for defense genes, while panel (b) presents the scores for general genes. Each panel compares the performance of context-dependent BERT embeddings and sequence-based ESM2 embeddings across different train/test splits: Random, Family-level, Order-level, and Class-level.

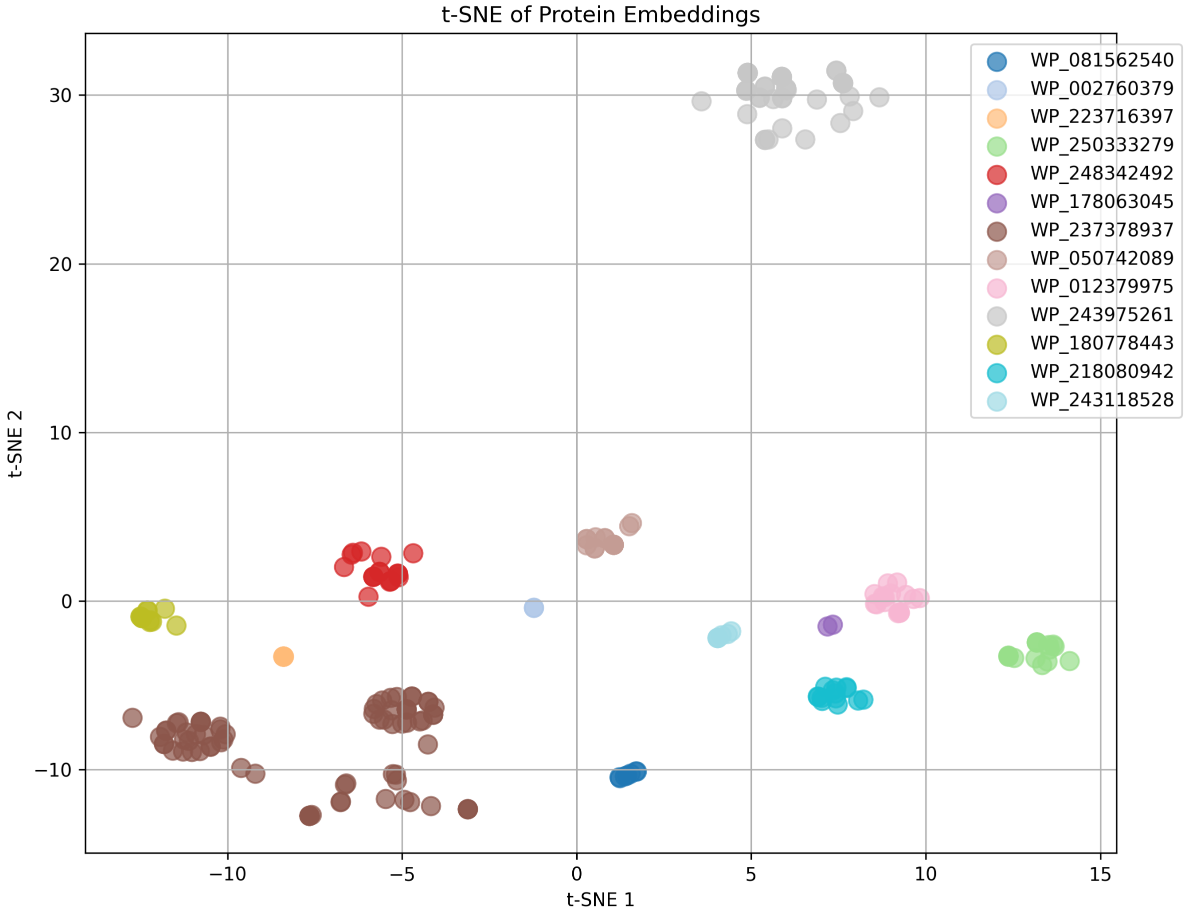

Figure 5: Visualization of Cas2 protein embeddings, serving as a case study. The legend displays various protein identifiers, all annotated with the InterPro ID IPR010152, which corresponds to Cas2 proteins. The plot reveals two distinct clusters: a main cluster and a smaller, remote cluster. This separation suggests that some Cas2 proteins exist in different genomic contexts. The distinct clustering highlights the importance of considering genomic context when analyzing protein function, even within the same protein family.

#### 2 Supplementary Tables

Table 1: Statistical analysis of defense proteins across taxonomic splits for BERT and ESM2 embeddings.

| | | BERT | | ESM2 | |
|---|---|---|---|---|---|
| Group1 | Group2 | p-adj | reject | p-adj | reject |
| Random | Family | 0.0 | **True** | 0.614 | False |
| Random | Order | 0.0 | **True** | 0.0.101 | False |
| Random | Class | 0.0 | **True** | 0.036 | **True** |
| Family | Order | 0.890 | False | 0.481 | False |
| Family | Class | 0.003 | **True** | 0.198 | False |
| Order | Class | 0.020 | **True** | 0.931 | False |

Table 2: Statistical analysis of general proteins across taxonomic splits for BERT and ESM2 embeddings.

| | | BERT | | ESM2 | |
|---|---|---|---|---|---|
| Group1 | Group2 | p-adj | reject | p-adj | reject |
| Random | Family | 0.0 | **True** | 0.121 | False |
| Random | Order | 0.0 | **True** | 0.172 | False |
| Random | Class | 0.0 | **True** | 0.286 | False |
| Family | Order | 0.040 | **True** | 0.995 | False |
| Family | Class | 0.002 | **True** | 0.928 | False |
| Order | Class | 0.569 | False | 0.982 | False |

Table 3: Class-specific performance of contextualized word2vec embeddings

| Class | Precision | Recall | F1-score |
|---|---|---|---|
| Amino sugar and nucleotide sugar metabolism | 0.606 | 0.699 | 0.649 |
| Benzoate degradation | 0.595 | 0.658 | 0.625 |
| Energy metabolism | 0.505 | 0.340 | 0.406 |
| Oxidative phosphorylation | 0.910 | 0.777 | 0.838 |
| Porphyrin and chlorophyll metabolism | 0.764 | 0.680 | 0.719 |
| Prokaryotic defense system | 0.741 | 0.799 | 0.769 |
| Ribosome | 0.819 | 0.777 | 0.797 |
| Secretion system | 0.787 | 0.738 | 0.762 |
| Two-component system | 0.611 | 0.703 | 0.654 |

Table 4: Class-specific performance of BERT embeddings

| Class | Precision | Recall | F1-score |
|---|---|---|---|
| Amino sugar and nucleotide sugar metabolism | 0.872 | 0.940 | 0.904 |
| Benzoate degradation | 0.851 | 0.886 | 0.868 |
| Energy metabolism | 0.816 | 0.444 | 0.575 |
| Oxidative phosphorylation | 0.958 | 0.876 | 0.915 |
| Porphyrin and chlorophyll metabolism | 0.922 | 0.910 | 0.916 |
| Prokaryotic defense system | 0.951 | 0.894 | 0.922 |
| Ribosome | 0.915 | 0.992 | 0.952 |
| Secretion system | 0.921 | 0.882 | 0.901 |
| Two-component system | 0.853 | 0.912 | 0.882 |

Table 5: Comparison of AUC Scores for BERT's Top 20 Biological Process GO Terms vs. DeepGO-SE

| | AUC BERT | AUC DeepGO-SE | Description |
|---|---|---|---|
| GO:0046181 | 1.000 | 0.500 | Ketogluconate catabolic process |
| GO:0000717 | 1.000 | 0.500 | Nucleotide-excision repair, DNA duplex unwinding |
| GO:0042150 | 1.000 | 0.500 | Plasmid recombination |
| GO:0006268 | 0.990 | 0.500 | DNA unwinding involved in DNA replication |
| GO:0031648 | 0.990 | 0.500 | Protein destabilization |
| GO:0006289 | 0.990 | 0.500 | Nucleotide-excision repair |
| GO:0048473 | 0.990 | 0.500 | D-methionine transmembrane transport |
| GO:0051454 | 0.990 | 0.496 | Intracellular pH elevation |
| GO:0070581 | 0.990 | 0.500 | Rolling circle DNA replication |
| GO:0019835 | 0.990 | 0.683 | Cytolysis |
| GO:0046417 | 0.990 | 0.842 | Chorismate metabolic process |
| GO:0032978 | 0.990 | 1.000 | Protein insertion into membrane from inner side |
| GO:0042867 | 0.990 | 0.500 | Pyruvate catabolic process |
| GO:0034258 | 0.990 | 0.500 | Nicotinamide riboside transport |
| GO:0002143 | 0.990 | 0.498 | tRNA wobble position uridine thiolation |
| GO:0015879 | 0.980 | 0.998 | Carnitine transport |
| GO:1902777 | 0.980 | 0.498 | 6-sulfoquinovose(1-) catabolic process |
| GO:1900232 | 0.980 | 0.993 | Negative regulation of single-species biofilm formation on inanimate substrat |
| GO:0015825 | 0.980 | 0.500 | L-serine transport |
| GO:0043487 | 0.980 | 0.653 | Regulation of RNA stability |

Table 6: Comparison of AUC Scores for BERT's Top 20 Molecular Function GO Terms vs. DeepGO-SE

| | AUC BERT | AUC DeepGO-SE | Description |
|---|---|---|---|
| GO:0008911 | 1.000 | 0.499 | Lactaldehyde dehydrogenase (NAD+) activity |
| GO:0050569 | 1.000 | 0.500 | Glycolaldehyde dehydrogenase (NAD+) activity |
| GO:0015386 | 1.000 | 0.489 | Potassium:proton antiporter activity |
| GO:0070573 | 1.000 | 0.499 | Metallodipeptidase activity |
| GO:0004333 | 1.000 | 0.498 | Fumarate hydratase activity |
| GO:0015385 | 1.000 | 0.479 | Sodium:proton antiporter activity |
| GO:0035527 | 1.000 | 0.500 | 3-hydroxypropionate dehydrogenase (NADP+) activity |
| GO:0080146 | 0.990 | 1.000 | L-cysteine desulfhydrase activity |
| GO:0008768 | 0.990 | 0.499 | UDP-sugar diphosphatase activity |
| GO:0071522 | 0.990 | 1.000 | Ureidoglycine aminohydrolase activity |
| GO:0034736 | 0.990 | 0.500 | Cholesterol O-acyltransferase activity |
| GO:1902670 | 0.990 | 1.000 | Carbon dioxide binding |
| GO:0015112 | 0.990 | 0.749 | Nitrate transmembrane transporter activity |
| GO:0070678 | 0.990 | 0.999 | Preprotein binding |
| GO:0097163 | 0.990 | 0.994 | Sulfur carrier activity |
| GO:0019172 | 0.990 | 0.750 | Glyoxalase III activity |
| GO:0047178 | 0.990 | 0.500 | Glycerophospholipid acyltransferase (CoA-dependent) activity |
| GO:0034618 | 0.980 | 0.500 | Arginine binding |
| GO:0050565 | 0.980 | 0.500 | Aerobactin synthase activity |
| GO:0015528 | 0.980 | 0.500 | Lactose:proton symporter activity |

Table 7: Comparison of AUC Scores for BERT's Top 15 Cellular Component GO Terms vs. DeepGO-SE

| | AUC BERT | AUC DeepGO-SE | Description |
|---|---|---|---|
| GO:0009317 | 1.000 | 0.998 | Acetyl-CoA carboxylase complex |
| GO:0045283 | 1.000 | 0.500 | Fumarate reductase complex |
| GO:1990586 | 1.000 | 0.500 | Divisome complex |
| GO:0009329 | 1.000 | 1.000 | Acetate CoA-transferase complex |
| GO:0009328 | 1.000 | 0.498 | Phenylalanine-tRNA ligase complex |
| GO:0045252 | 1.000 | 0.500 | Oxoglutarate dehydrogenase complex |
| GO:0033281 | 1.000 | 1.000 | TAT protein transport complex |
| GO:0017117 | 1.000 | 0.500 | Single-stranded DNA-dependent ATP-dependent DNA helicase complex |
| GO:0022625 | 0.990 | 0.690 | Cytosolic large ribosomal subunit |
| GO:0009358 | 0.990 | 0.500 | Polyphosphate kinase complex |
| GO:0009419 | 0.990 | 0.750 | Pilus tip |
| GO:0000796 | 0.990 | 0.500 | Condensin complex |
| GO:0009898 | 0.990 | 0.804 | Cytoplasmic side of plasma membrane |
| GO:0009350 | 0.990 | 0.500 | Ethanolamine ammonia-lyase complex |
| GO:0045261 | 0.980 | 0.494 | Proton-transporting ATP synthase complex, catalytic core F(1) |
